# Balancing precision with plasticity: Redefining the roles of transcription factors in early cell fate specification

**DOI:** 10.1101/2025.02.11.636561

**Authors:** Shiva Abbasi, Sijie Li, George Charitos, Evangelia Chrysostomou, Wenjiang Zhou, Panagiotis Apostolidis, Priyanshi Borad, Peng He, Theodora Koromila

## Abstract

Early embryogenesis is governed by intricate gene regulatory networks that orchestrate cell fate specification across all three germ layers. While the role of transcription factors (TFs) and pioneer factors (PFs) in shaping chromatin accessibility is well established, the mechanisms coordinating spatiotemporal gene expression before gastrulation remain poorly understood. Here, we investigate how a coordinated network of early-acting regulators, including the pioneer factors Zelda (Zld), GAGA factor (GAF), and the pioneer-like timing factor Odd-paired (Opa/ZIC3), along with Dorsal (Dl), Twist (Twi), the Polycomb subunit Enhancer of Zeste (E(z)), and the co-activator CBP, govern early transcriptional events in the *Drosophila* embryo.

Using integrated chromatin accessibility, histone modification, TF binding, and transcriptomic data (bulk and single-cell), we identify two distinct classes of regulatory elements that govern early gene expression. The first class includes distal, PF-dependent enhancers that are inaccessible early and require PFs such as Opa and Zld to overcome Su(H) activities and H3K27me3-mediated repression and initiate transcription. The second class, enriched for PcG domains, comprises proximal enhancers that are accessible from the earliest nuclear cycles, despite being marked by H3K27me3 and bound by E(z). These regions are preloaded with RNA Polymerase II and positioned near promoters, suggesting a regulatory state primed for activation. While bound by multiple early TFs, these enhancers maintain accessibility independently of PF activity, functioning instead as autoregulatory elements that rely on PFs for maintenance rather than initiation.

This study expands our understanding of how early TFs and PFs coordinate to balance transcriptional precision and developmental plasticity across germ layers, and redefines the role of PFs as both activators and stabilizers of gene regulatory architecture. Within this framework, Opa regulates key mesodermal and neuroectodermal genes such as *slp1* and *odd*, and its activity is modulated by Suppressor of Hairless (Su(H)). At earlier stages, Su(H) represses Opa target genes, but rising Opa levels at later timepoints override this repression, enabling activation.

Together, these findings support a dual-mode enhancer logic: one mode driven by PF-mediated chromatin opening at distal, repressed enhancers, and another where proximal, pre-accessible enhancers within PcG-marked regions are reinforced by PF binding. This study redefines the role of PFs in early development, emphasizing their maintenance and stabilizing roles at poised enhancers, and provides a framework for how embryos balance transcriptional plasticity with spatial precision across all germ layers.

## Introduction

In all animals, embryonic cells undergo specific signaling dynamics that determine the various cell fates and develop different complex tissues^1^. Interactions among transcription factors (TFs) and cis-regulatory elements (CREs) of their downstream genes establish a complex network that governs cell differentiation and specification^2^. Elucidating this transcriptional network will enhance our understanding of the intricate mechanisms underlying cell-specific differentiation. In several studies aimed at understanding the interactions between sequence-specific TFs and their enhancers and how they are regulated in embryogenesis, *Drosophila* has been found to be the ideal experimental model^3^. *Drosophila*’s heart genes have been found to share homology with equivalently functional vertebrate heart gene^4^. Moreover, strong evidence derives from the fact that studies in congenital heart disease (CHD) use *Drosophila* as their main model system^5^. Understanding how the interactions between regulatory elements govern the spatiotemporal expression of key cardiac genes could reveal critical pathways that, when dysregulated, contribute to the development of specific heart diseases. *Drosophila’s* heart originates from the embryonic mesoderm^6^, and the initial phases of heart formation follow remarkably similar patterns across species, from flies to complex mammals such as humans^7^. These similarities encompass the origin of cardiac precursor cells, structural organization, regulatory networks, and how the cardiac progenitors migrate towards the middle of the embryo^4,8^.

In the past decades, the researchers have shed light on the functional and regulatory roles of pioneer factors (PFs) that have the property to bind and open the chromatin at specific enhancer regions, so that other TFs can bind and activate zygotic gene expression^9–11^. TFs such as Dorsal (Dl, orthologous to NF-κB)^12^, Twist (Twi, orthologous to human twist family bHLH TF 1)^13^, Odd-paired (Opa, orthologous to zinc finger protein of the cerebellum 3 (ZIC3))^14^, Suppressor of hairless (Su(H), orthologous to human recombination signal binding protein for immunoglobulin kappa J region (RBPJ))^15^ are present in *Drosophila* embryo. Dl is an early expressed TF required for dorsal-ventral patterning of the embryo^16^. Twi is one of the earliest expressed zygotic genes and a crucial mesodermal TF, which is necessary for the dorsal-ventral pattern formation in *Drosophila* embryos activated by maternally expressed Dl in the ventral part of the embryo^17,18^. Opa acts as a timing factor and is expressed uniformly throughout the trunk region of the embryo^19,20^. Mutations in Opa homologue in mammals, ZIC3, have been identified in congenital heart disease^21^. Su(H) is a critical TF that interacts with the evolutionary conserved Notch signaling pathway, which is involved in heart cells specification. In cardioblast cells, through Su(H), Notch signaling represses the enhancer of pericardial cells^22^. Su(H) contribute to stage-specific regulation and has a repressive role during cellularization and regulating transcriptional levels and counterbalance to Dl and Zelda (Zld) activating roles^23–25^. GAF encoded by the Trithorax-like (Trl) gene, functions as a pioneer transcription factor during the onset of zygotic genome activation. GAF plays a crucial role in initiating gene expression during early *Drosophila* development^26^. Opa is involved in the temporal and concentration-dependent regulation of several pair-rule and segment-polarity genes, including *even-skipped* (*eve*), *engrailed* (*en*), *paired* (*prd*), *fushi tarazu* (*ftz*), and *odd-skipped* (*odd*)^19^. Another notable regulatory role of Opa is that it involves in frequency doubling by controlling the accessibility of regulatory regions^9^. Based on their regulatory elements and expression pattern during early cellularization, pair-rule genes can be classified into two categories: primary and secondary^19,27^. Primary pair-rule genes, such as *odd* and *runt* (*run*) are regulated by stripe-specific elements^27^, and secondary genes such as *sloppy_paired* (*slp)* regulating by one anterior stripe-specific element and a zebra element regulating their expression in the middle part of the embryo^27^. It is important here to mention that regulatory targets (e.g., *prd, odd, slp, run*) respond to different Opa concentration levels depending on a varying threshold of Opa’s activity that needs to be reached for full target expression during gastrulation. Lower levels of Opa result in delayed events in gastrulation causing, for instance, the *slp* stripes to appear slightly delayed in embryos with weak Opa expression^19^.

Opa acts as a timing factor, regulating the transition from the 7-stripe to the 14-stripe expression pattern of pair-rule TFs is not observed in *opa* mutant embryos during gastrulation. At the onset of gastrulation, changes in pair-rule genes has been reported in *opa* mutants^19^. In this study, we performed Fluorescence *In Situ* Hybridization (FISH), used publicly available datasets, including assay for transposase-accessible chromatin with sequencing (ATAC-seq), and chromatin immunoprecipitation (ChIP) assays with sequencing (ChIP-seq) for Dl, Opa, Su(H), GAF, and Twi^11,23,28,29^, as well as single-cell^30^ and bulk RNA-seq^11^ of *Drosophila* embryo. Our primary objective was investigating the role of Opa in the regulation of pair-rule genes during pre-gastrulation, with a particular focus on its involvement in early cardiac cell specification. Our results support the hypothesis that Opa plays a role in early gene expression, specifically by regulating two genes in cardiac gene regulatory network, *slp1* and *odd* before gastrulation^31,32^. While our previous studies have demonstrated the role of Opa in brain cell specification at this developmental timepoint, our findings here reveal that Opa also regulates cardiac-specific gene expression before gastrulation^33^. The regulatory network governing embryonic gene expression is highly complex and dynamic across various developmental stages, including early mesoderm formation and later stages of cardiac differentiation^34–36^. While much attention has been given to gene regulation at later timepoints, our study aims to provide insight into the primordial regulatory events that contribute to cardiac cell specification. The results reveal two enhancer modes in early embryogenesis: one that depends on pioneer factors to open chromatin and drive context-specific transcription, and another that is accessible early and maintained by promoter proximity and Pol II, with pioneer factors only reinforcing activity. Together, they provide a flexible yet stable system for precise gene regulation.

## Results

### Opa’s regulatory role in embryonic development before gastrulation

There are well-studied cis-regulatory elements identified in the *slp1* genomic region. The distal early stripe element (*DESE*)^37,38^, is regulating 14-stripes of *slp1* at the end of nc14D. Studies show that Opa is required for *DESE* activation at late nc14^11^. *in situ* hybridization of *slp1* shows that in control embryos, late *slp1* stripes appear at nc14D. However, in *opa* knockdown embryos, these late stripes are absent (Fig. 1A), suggesting that *DESE* is an Opa-dependent late stripe enhancer, as we see loss of *DESE* stripes in *opa* knockdown embryos at nc14D. The JASPAR binding sites of Dl, Twi, Opa, Su(H), and GAF were explored within the *DESE* enhancer (Fig. 1G, H). We also examined *prd-01, odd _basal-1, run_2.6B, and sna_prox 2.2* regulatory regions. Based on the TFs binding profiles, we categorized Opa target regulatory elements into two groups: group I (Fig. 1I), which includes distal enhancers bound by Opa but not GAF, and group II, which includes proximal enhancer co-bound by both Opa and GAF. *slp1 DESE* and prd_01 are classified as group I, while *odd-basal-1*, *run_2.6B*, and *sna_prox 2.2* fall into group II. *prd*_*01* showed similar pattern as *DESE* genomic region in *slp1* locus. We categorized these loci as group I, which Opa impact them without GAF.

**Fig. 1.**
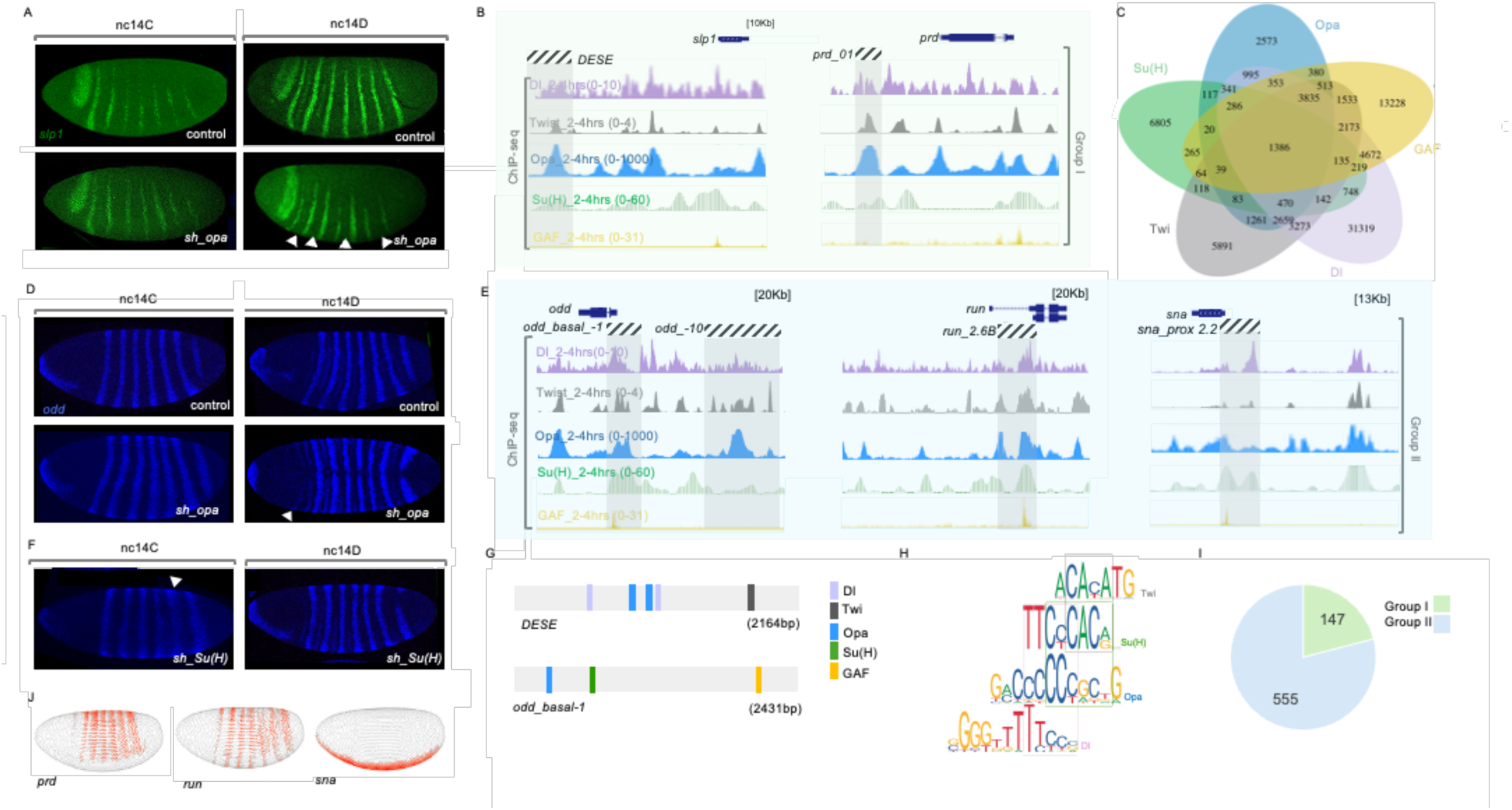
Opa regulates mesodermal genes through two distinct mechanisms: direct and indirect. A) *slp1* transcriptional dynamics before gastrulation (stage 5 late). In situ hybridization using riboprobes to *slp1* of wild-type (yw for nc14C and UAS-lacZ.Tub.Gal4 for nc14D) and sh_opa embryos at stage nc14C (top) and nc14D (bottom). Altered *slp1* expression patterns were observed in *sh_opa* embryos, with the loss of stripes indicated by white arrows in the figure (¾ embryos). B, E) ChIP-seq of Dl at 2-4hrs (lavender), Twi at 2-4hrs (gray), Opa 2-4hrs (blue) and Su(H) at 2-4hrs (green), and GAF (yellow) from available data10,22,26,27 in group I genes, slp1 and prd (B) and *odd*, *run* and *sna* in group II (E). All the regulatory elements are highlighted in gray. C) Venn diagram created using publicly available ChIP-seq data showing overlapping and unique binding sites among, Dl (lavender)Twi (gray) and Opa (blue), GAF (yellow) and Su(H) (green)10,22,27, peak calling data of TFs without any filtering were used. D) Opa’s contribution to the regulation of odd gene expression and patterning during the nc14C and nc14D stages. FISH staining performed using odd riboprobe on control (wt) and sh-opa embryos at nc14C and nc14D. Arrowhead showing the altered phenotype in *sh_opa* embryos (½ embryos). F) In situ hybridization of *odd* in blue control (yw) and *sh_Su(H)* embryos, demonstrating the expression pattern *odd* at nc14C and nc14D. Arrowhead showing the changes in *odd* expression (3/4 embryos). G) Schematic representation showing the binding sites of the TFs (Dl, Opa, Su(H), GAF) in the genomic location of the odd_basal-1 (2431bp) and DESE (2164bp). H) JASPAR consensus binding sites for Su(H), Twi, Opa and DI. Each sequence logo represents a specific DNA binding site pattern. The height of each letter indicates the relative frequency of nucleotides (A, T, C, G) at each position, with taller letters indicating higher conservation. I) Pie chart showing the proportion of annotated regions in group I (n=147) and group II (n=555) at Zld bound regions (See supplemental 1.2F). J) Endogenous expression of *prd*, *run*, *sna* obtained from DVEX (Drosophila Virtual Expression eXplorer) database at stage 6.

We further hypothesized that Opa regulates *odd* through an indirect mechanism before gastrulation, since we noticed that *odd* expression shows expedited phenotype when Opa is knocked down (Fig. 1D). At nc14C, low Opa levels allow Su(H) to repress Opa-mediated activation of *odd*. However, by nc14D, as Opa levels increase, Opa becomes more dominant in regulating *odd* expression. This is evident at nc14D, where a phenotype change is observed in *sh_opa* embryos, corresponding to sufficient Opa levels. Specifically, in the absence of Opa at nc14D, the repressor Su(H) can’t sufficiently reduce *odd* expression levels, resulting in an accelerated *odd* expression pattern that reflects a later-stage phenotype (Fig. 1D and supplementary fig. 1.1E). Opa is a well-established activator, and as such, we would expect *odd* expression to be downregulated or remain at an earlier developmental stage. However, the observations here deviate from this expectation, leading us to propose that Opa exerts an indirect regulatory role in modulating *odd* expression through the action of a repressor. However, further studies are needed to explore other target genes of Opa and to further develop this hypothesis. Motif analysis (Twi, Su(H), Opa and Dl JASPAR shows that Dl, Twi, Opa, Su(H) bind to odd regulatory regions (Fig. 1G, supplementary fig. 1.1F). Thus, we suggest that at later stages, when Opa levels are higher, Opa regulates odd, but earlier Su(H) facilitates odd repression. To further validate our findings, we performed in situ hybridization of slp1 and odd staining on sh-Su(H) embryos. As expected, in Su(H) knockdown embryos, we observed increased odd expression levels, with wider expression stripes at nc14C (Fig. 1F). At the later stage, nc14D, when Opa’s levels are elevated, Opa acts as an activator and overcomes Su(H) effect. These results support our hypothesis that Su(H) and Opa mutually regulate each other’s expression during early stages, with Su(H) acting as a repressor. However, as Opa levels increase, it appears to act as an activator, suggesting a stage-specific balance in their regulatory relationship.

From our analysis *Su(H)* knockdown leads to impaired *odd* and *slp1* expression and morphological defects. In situ hybridization of *odd* and *slp1* in *sh_Su(H)* embryos revealed alterations in the expression of these genes. During cellularization at nc14B/C, we observed impaired expressions of both *odd* and *slp1* genes (Supplementary fig. 1F). Previous studies have shown that ectopic expression of Su(H) leads to defects in cellularization and alterations in gene expression23. In our study, we observed aberrant expression patterns in *sh_Su(H)* embryos during cellularization, indicating that loss of Su(H) function not only disrupts normal gene regulation but it is also responsible for morphological abnormalities.

We studied *odd_basal_-1*^27^ regulatory enhancer that support 7-stripe expression of *odd* at early cellularization. The occupancy of Dl, Twi, and GAF at these regions indicates active regulatory engagement, supporting the notion that these enhancers are functional from the earliest stages of embryogenesis. The occupancy of Dl, Twi, and GAF at these regions indicates active regulatory engagement, supporting the notion that these enhancers are functional from the earliest stages of embryogenesis.Similarly, Opa and Su(H) bind to these regulatory regions (Fig. 1E), supporting the hypothesis of coregulation by these two TFs. To further explore the discussed mechanism and identify additional examples of this regulatory logic across the genome, we analyzed the binding profiles of Dl, Twi, Su(H), Opa, and GAF. This analysis revealed 170 loci in group I and 1,386 loci in group II (Fig. 1C). Among these, 231peaks in group I, and 655 peaks in group II are located within 100 bp of REDfly^39–44^ cis-regulatory modules (Supplementary Fig. 1.2A). Incorporating Zld into the analysis identified 200 group I regions and 672 group II regions, of which 147 and 555 peaks, respectively, overlapped with REDfly enhancers (Supplementary fig. 1.2C, D). Both groups include regions within enhancers and promoters (Supplementary fig. 1.2B). The significant overlap of both group I and group II peaks with REDfly cis-regulatory modules supports the idea that these co-bound regions represent functional enhancers active during development, rather than random TF-binding events.

Chromatin accessibility plays a critical role in facilitating transcription factor recruitment to regulatory regions and maintaining dynamic control of gene expression during development^45,46^. In *odd_-10* region chromatin remained unchanged upon overexpression of Opa. In *odd_basal_-1* region, however, chromatin is unchanged upon overexpression and *sh_opa* (Fig. 2A). Group II, which represents the genomic locus bound by both Opa and GAF, includes *odd_basal_-1, run_2.6B*, supporting striped pattern of *run*^47^, (Supplemental fig. 2.1A). Although Dl occupies enhancers in both groups, it does not appear to regulate chromatin accessibility, as loss of Dl (gd^7^) does not alter chromatin occupancy at these loci at the examined timepoints.

**Fig. 2.**
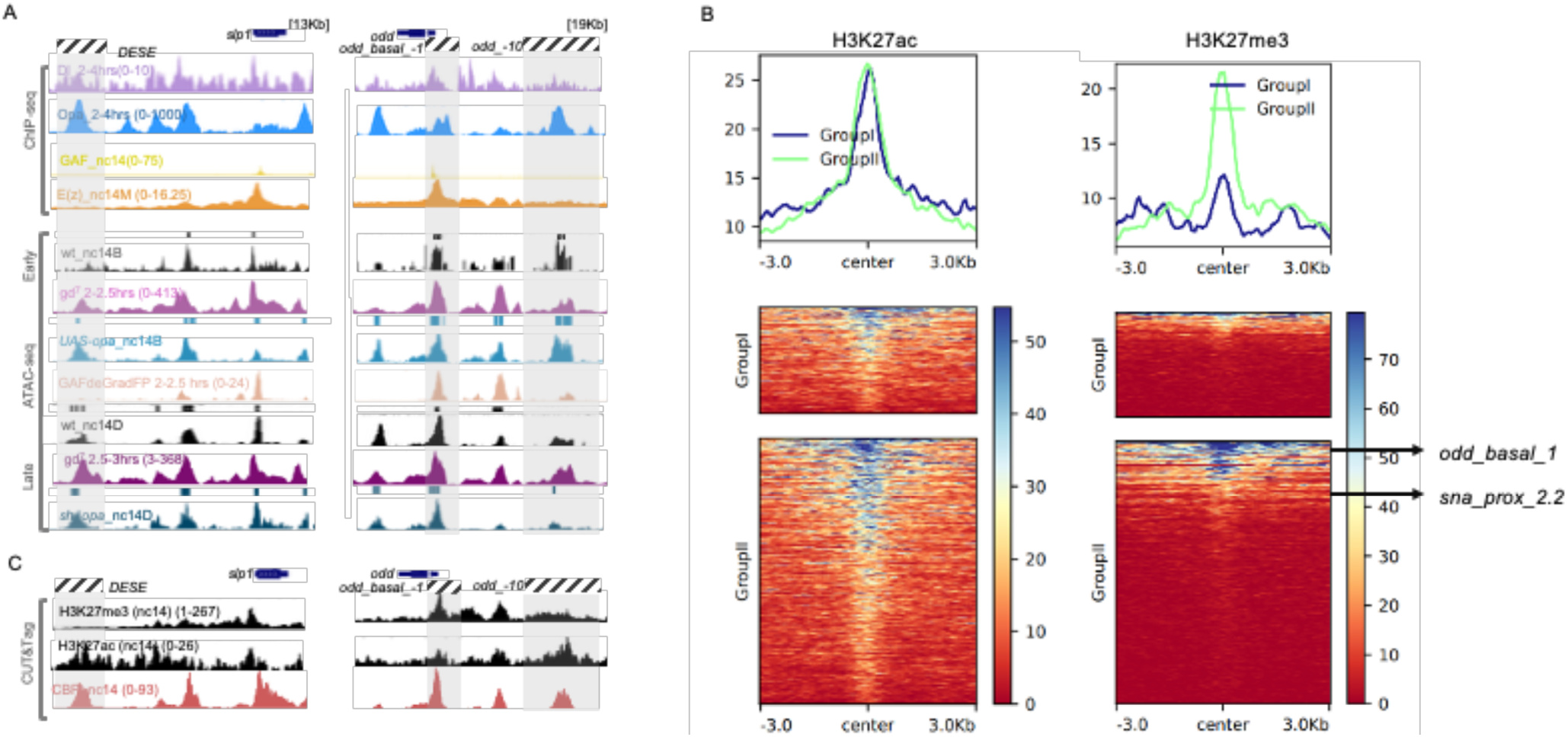
Transcription factor binding, chromatin accessibility, and functional enrichment distinguish group I and group II genes. A) ChIP-seq of Dl at 2-4hrs (lavender), Twi at 2-4hrs (gray), Opa 2-4hrs (blue) and Su(H) at 2-4hrs (green), and GAF (in yellow) from available data^11,23,28,29^ in *slp1* (group I) and *odd* (group II). Genome browser tracks representing the chromatin accessibility (ATAC-seq) wildtype control for nc14B (wt) (top black), *UAS-opa*_nc14B (light blue), wt control for nc14D (bottom black), *sh_opa* knockdown at nc14D (dark blue), GAF degraded embryos (2–2.5 hrs), Dl mutant/embryos lacking nuclear Dorsal (2-2.5hrs) and gd^7^/embryos lacking nuclear Dorsal (2.5-3 hrs). B) Heatmaps were generated to visualize the signal intensities of histone modifications H3K27ac, H3K27me3, centered on genomic regions corresponding to group I and group II. C) Occupancy profiling (Available data of Cut&Tag data of H3K27me3 and H3K27ac histone marks (nc14) and CBP at nc14 in *slp1* and *odd_basal* genes.

To explore the epigenomic landscape of the two Opa-target genes groups, we tested the two histone modification that marks active chromatin and transcriptional silencing, H3K27ac and H3K27me3, respectively^48^. These histone marks have been used to study chromatin landscape in *Drosophila* embryo in previous studies ^49–51^. Our analysis using the published histone marks data^49^ indicates that H3K27ac shows similar enrichment at group I (*slp1*, *prd*) and group II (*odd*, *run*, *sna*) gene loci, whereas H3K27me3 is absent at group I and present at group II, suggesting the ability of PFs to open the chromatin, when chromatin is closed from nc11 to 13 (Supplementary fig. 2.1B, 2.3A). H3k27me3 show higher occupancy in group II loci across genome, while H3K27ac association showed no significant differences between the two groups (Fig. 2B). Polycomb Repressive Complex 2 (PRC2) contains the Enhancer of Zeste (E(z)) subunit, which functions as a methyltransferase and establishes a transcriptionally silent chromatin state necessary for proper gene regulation^52^. To directly assess the role of Polycomb-mediated repression at these loci, we next examined E(z) occupancy using ChIP-seq at mid nc14^53^. This allowed us to determine whether the differential H3K27me3 enrichment observed between two groups correlates with E(z) binding. (Fig. 2A, supplementary fig. 2.1A). E(z) binds to group II genes. To study the association of the two groups with PcG domains, we further compared them with previously defined Polycomb group (PcG) domains^53^ to evaluate how these pioneer factor-targeted loci relate to broader repressive chromatin landscapes. Genes associated with group II regions that overlap with PcG domains are more strongly linked to mesoderm and cardioblast differentiation, whereas genes in non-PcG regions are more enriched for neuronal differentiation. Both groups, however, share cell differentiation as their primary function Supplementary fig. 2.1D, supplementary data 1, 2).

We examined the genomic regions where group IIB enhancers (Opa, GAF, Twi, Dl, Zld) overlap with E(z) peaks, as well as with accessible regions from the Dl mutant, Opa ectopic expression, and Opa knockdown datasets, 1,470 REDfly enhancers identified between group II proximal regions (±2 kb from the TSS) and PcG domains (Fig. 5D). We then analyzed the functions of the genes in different groups. Group I show function more enriched in cell fate commitment and morphogenesis, while group IIB cell fate and transcription regulation (Fig. 5C).

### Proximal autoregulatory feedback enhancers are not regulated by PFs

Cis-regulatory regions have been shown to play a critical role in the autoregulatory feedback control of gene expression. In particular, the *sna* gene exhibits negative autoregulation mediated through its proximal enhancer^54^. This autoregulatory mechanism appears to function by modulating the input of general activators, such as Dl^54^. Autoregulatory feedback becomes established at these enhancers, allowing the *sna* locus to maintain its own transcriptional output. Notably, these autoregulatory enhancers exhibit autonomous activity and are not regulated by PFs. They preserve both chromatin accessibility and transcriptional activity independently of canonical PFs such as Opa, despite Opa’s binding to these regions and concurrent developmental activity. Interestingly, chromatin accessibility at these regulatory enhancers is not significantly altered by the presence or absence of PFs, indicating a PFs-independent mechanism of enhancer maintenance (Fig. 2, supplementary fig. 2.1, 2.2). This regulatory logic is conserved along both the DV and anteroposterior (AP) axes, suggesting a shared regulatory architecture across spatial patterning domains.

Two distinct classes of enhancers can be distinguished: autoregulatory feedback enhancers that function independently of known PFs (group II) and maintain activity autonomously, and Opa-dependent enhancers that require Opa for both accessibility and activation, highlighting a context-dependent enhancer logic during mesodermal patterning (group I).

These mechanisms in group I and II are not restricted to a single germ layer, but operates across and between multiple layers. For example, the gene *odd* serves as a marker expressed in both mesoderm and endoderm. Likewise, *sna* is a mesoderm marker. These examples show that the mechanisms governing enhancer-driven expression are not confined to one lineage but instead enable the coordinated regulation of gene expression across multiple embryonic cell types. This view offers a unified model for understanding the chromatin-mediated control of cell fate, developmental plasticity, and germ layer specification in early embryos.

### TFs dynamics regulate gene expression patterns but not expression levels during cellularization in *Drosophila* embryos

Gene expression is highly dynamic during the early stages of embryogenesis. In this study, we chose to focus on both primary pair-rule genes, e.g. *odd,* and secondary pair-rule genes, e.g. *slp1,* as well as Dorsal Ventral (DV) mesoderm-specific genes, e.g. *sna* to investigate the role of Opa in mesodermal patterning before gastrulation. The genes are expressed early in development, with *odd* forming stripes in the trunk region of the embryo, while *slp1* is restricted to anterior expression. These initial patterns evolve into stripe formations during cellularization (Supplementary fig. 1E). Additionally, Opa exhibits expression in the trunk region at mid-cellularization (Supplementary fig. 1.1A). Previous studies have demonstrated that Opa functions as a PF in cell specification, initiating its role at cellularization before gastrulation begins^33^. To investigate the broader role of Opa on cardiac gene regulation in *Drosophila* embryogenesis, we compared gene expression profiles between wild-type (*yw*) and *opa* knockdown (MTD-Gal4, *opa* short hairpin (sh) RNAi (*sh_opa*) embryos at nuclear cycle (nc) 14B, 14C and 14D, using fluorescent hybridization and single-cell^55^ and bulk RNA sequencing (RNA-seq) data from previous studies^11^. Our RNA-seq analysis showed that in *opa* knockdown embryos expression of the majority of mesoderm specific genes have not been significantly changed (Fig. 3C). Specifically, we observed significant changes in the expression levels of *odd* and *slp1* between nc14A and nc14C, as well as nc14A and nc14D (Supplementary fig. 1.1C). Despite the known dynamic nature of the expression patterns of these genes during these timepoints (Supplementary fig. 1.1D) no significant difference in either *slp1* or *odd*’s expression levels were detected between nc14C and nc14D which could possibly explain the lack of significant changes in the absence of the timing factor Opa at single-embryo RNA-seq data at nc14D.

**Fig. 3.**
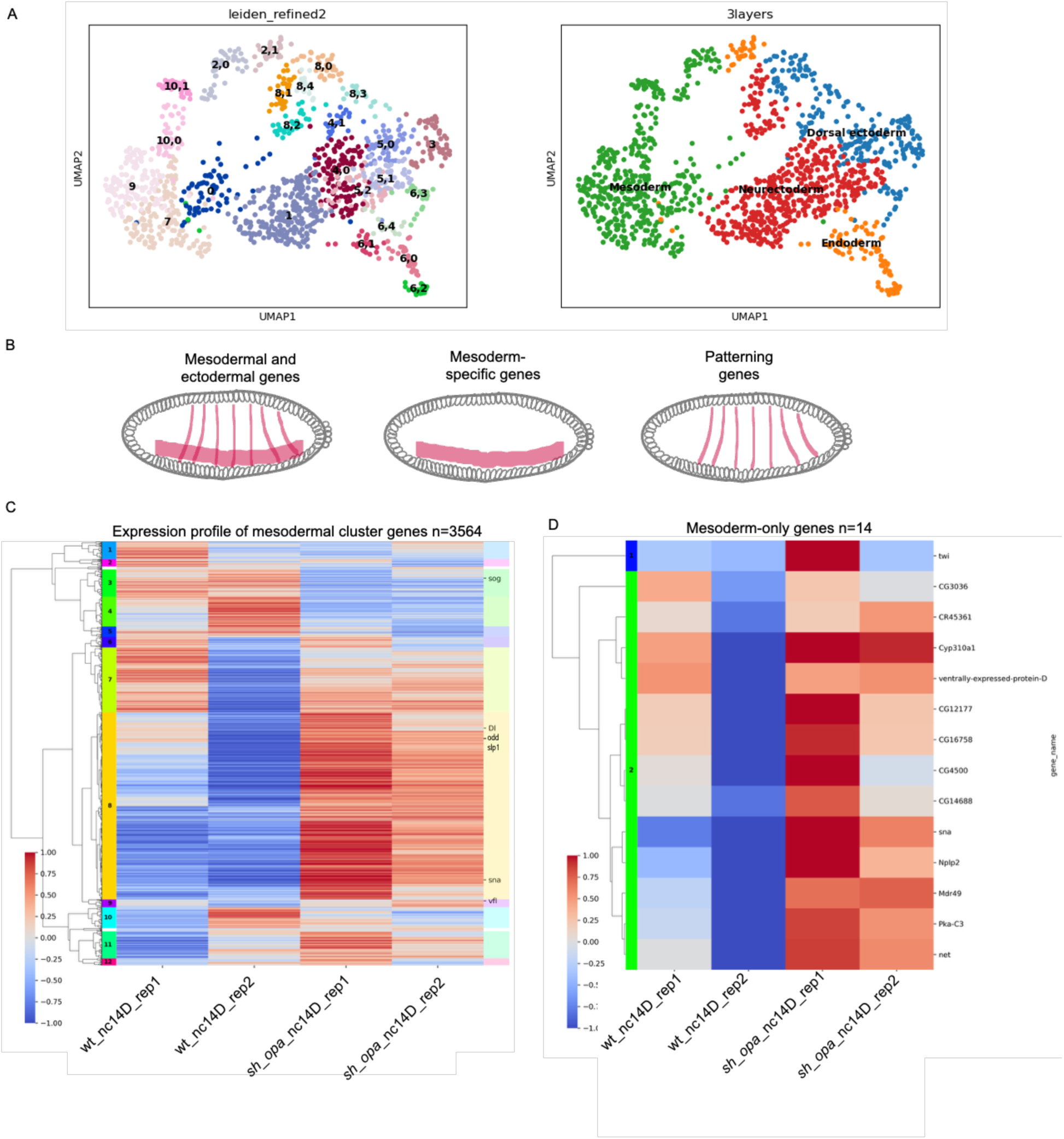
Single-cell and bulk RNASeq integration reveals Opa-dependent gene expression. A)Visualization of single-cell RNA-seq clustering (left) and germ layers (mesoderm, neuroectoderm, dorsal ectoderm, and endoderm) identification (right). B) Schematic representation of mesodermal cluster, mesoderm-specific, and mesodermal patterning genes in the *Drosophila* embryo. C) Heatmap illustrating expression profile of mesodermal cluster genes (n=3564), defined as the intersection between slightly filtered bulk RNA-seq data (retaining genes with TPM>10 in at least 2 samples) and genes from mesodermal clusters in the single-cell data. All heatmaps show centralized log2(TPM+1) values, with two biological replicates per condition (control and sh_opa). D) Heatmap of mesoderm-only genes within the filtered bulk RNA-seq dataset (retaining genes with TP >7 in at least 2 samples). All heatmaps show centralized TPM values, with two biological replicates per condition (control and *sh_opa*).

### Comparing single-cell transcriptome with RNA-seq of *Drosophila* embryo

Integrating single-cell RNA-seq data with bulk RNA-seq allows high-resolution analysis of gene expression changes in specific cell populations while harnessing the sensitivity and broad transcriptome coverage provided by bulk approaches. This combined strategy reveals subtle biological insights that might be missed using either method alone. We sought to identify differentially expressed genes in mesodermal cluster in *sh_opa* bulk RNA-seq data^11^ correlated with mesodermal clusters in single cell resolution expressions^30^. We used markers from different germ layers to cluster cells with similar expression profiles (Fig. 3A, supplementary fig. 3.2A-D, 3.3A,B).

This methodology helps us to detect the differentially expressed genes in such as *odd*, *slp1*, *Dl*, *sna* (upregulated) in mesodermal clusters (Fig. 3C, supplementary fig. 3A).

Enrichment of H3K27ac, a marker of active enhancers and promoters, suggest increased transcriptional activity in these regions. However, the lower levels of enrichment of H3K27me3 at gene promoters indicates a reduced repressive chromatin state, further supporting an active transcriptional landscape within these mesodermal clusters. These histone marks show similar patterns across different clusters (Supplementary fig. 3.1B)

Next, we decided to explore the mesodermal gene expression in different Opa-bound categories (Supplementary data 3, 4), groups I and II. Opa+/GAF-group exhibits distinct regulatory characteristics. Opa target genes lacking GAF binding are more frequently associated with non-mesodermal clusters (neuroectoderm and dorsal ectoderm)(Fig. 4A, B). Notably, *odd*, and *sna* found to be expressed in the majority of the mesodermal clusters (Fig. 4C, D) suggesting that the co-binding of Opa and GAF may be critical for the regulation of mesodermal gene expression, whereas Opa alone appears to play a broader role beyond mesodermal lineage specification. The *short gastrulation (sog*) gene is reported to lose expression in the absence of Opa^11^, which is consistent with our analysis showing that sog is downregulated under *opa* knockdown conditions (Fig. 4D). These differentially expressed genes are active in multiple tissues, and by integrating bulk and single-cell expression data, we can observe their tissue-specific expression and the differences in expression levels.

**Fig. 4.**
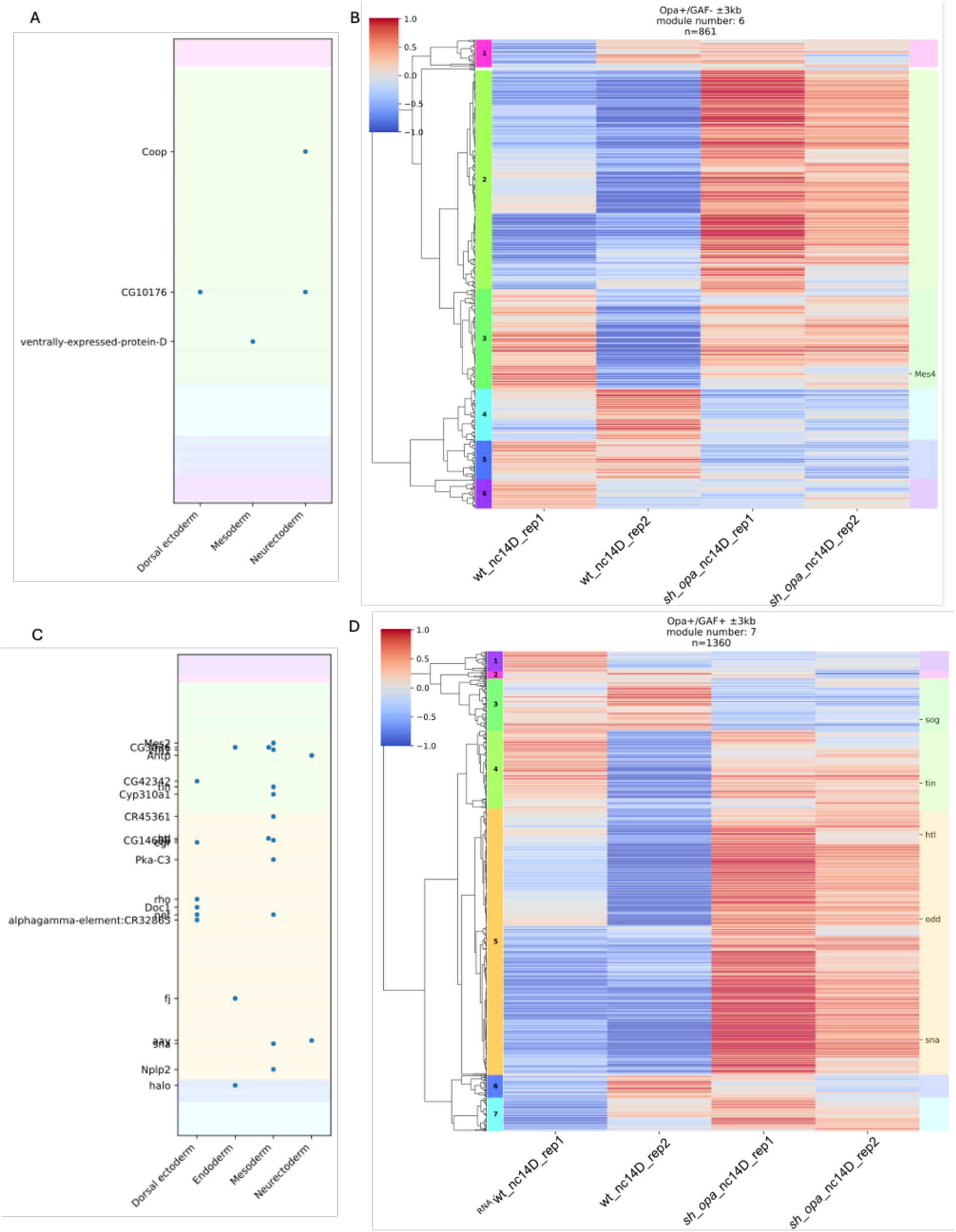
Bulk and single-cell integration of Opa/GAF regulation. Genes from slightly filtered bulk RNA-seq data are divided into Opa+/GAF− (B)Opa+/GAF+ (D) and. Both heatmaps show centralized log2 (TPM+1) values, with two biological replicates per condition (control *and sh_opa*). (A, B) Integration of single-cell data with whole-embryo transcriptome clustering. Y-axis: genes ordered as in the right heatmap; X-axis: cell type. A point indicates the expression of a cluster marker gene, and colors denote gene modules obtained from hierarchical clustering in C and D.

**Fig. 5.**
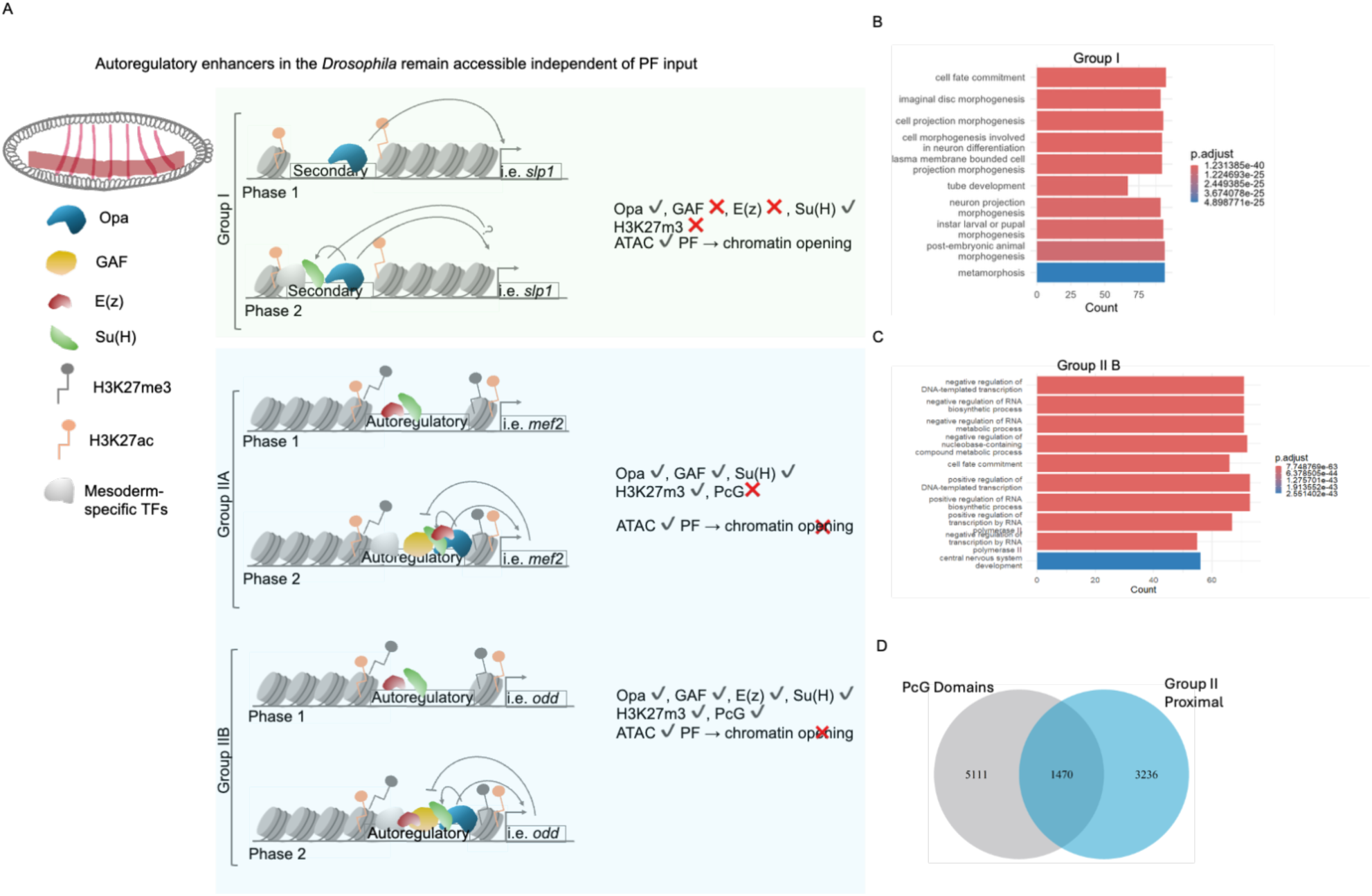
Dual enhancer logic in early embryogenesis. A) PF-dependent enhancers (group I) require pioneer factors to open chromatin and initiate transcription in a stage– and context-specific way. PF-independent, autoregulatory enhancers (group IIB) are accessible from early stages, maintained through promoter proximity and Pol II preloading, and strengthened, but not initially opened, by pioneer factor binding. B) Go analysis showing the biological function for group I, using ChIPseq data for Opa, Twi, Dl, Zld. C) Biological function of group IIB (Opa, GAF, Dl, Twi, Zld,) genes, shown in overlap with E(z), ATACseq peaks at gd^7^ (2-2.5)), gd^7^(2.5-3), UAS_Opa_nc14b, sh_Opa_nc14d, and PcG domains.

## Discussion

Our study challenges the prevailing paradigm of chromatin accessibility regulation by revealing a novel class of autoregulatory feedback enhancers in the *Drosophila* mesoderm that operate independently of canonical PF activity. We show that these enhancers remain accessible and transcriptionally active in the absence of functional input from factors such as Zld, GAF, and Opa, all of which are classically defined as pioneers of chromatin opening during early development. This finding reveals a new layer of regulatory logic in which enhancer accessibility and activity are not necessarily linked to pioneer factor engagement, but instead rely on intrinsic autoregulatory feedback loops. We propose that these enhancers serve as “mentors of the genome”: autonomous regulatory hubs that maintain accessibility and reinforce cell identity independently of upstream activation cues.

Using a combination of ATAC-seq, ChIP-seq, and single-nucleus multiomic profiling, we identified two mechanistically distinct classes of enhancers in the early embryo. One class is Opa-dependent, requiring Opa for chromatin accessibility and transcriptional activation (group I). The other class, encompassing autoregulatory feedback enhancers (group II), is both accessible and transcriptionally active despite being bound by, but functionally independent of, pioneer factors including Opa, Zld, and GAF. Strikingly, these PF-independent enhancers are enriched for the repressive histone mark H3K27me3, suggesting they operate within a chromatin landscape that is traditionally considered restrictive.

The decoupling of pioneer factor binding and chromatin accessibility highlights context-specific mechanisms in the regulation of mesodermal gene expression. While Opa and other pioneer factors regulate chromatin accessibility at many developmental loci, their binding at group II enhancers appears non-causative. Instead, these enhancers maintain accessibility and functionality through transcriptional autoregulation, possibly facilitated by persistent transcription factor occupancy and paused RNA Polymerase II (RNAPII), even in the absence of productive transcription. Overall, our results propose a dual-mode enhancer logic across the germ layers (Figure 5A):

1. PF-dependent enhancers (group I), which require pioneer factor activity to open chromatin and activate transcription in a stage– and context-dependent manner.
2. PF-independent, proximal autoregulatory enhancers (group II), which are accessible early, maintained by promoter proximity and Pol II preloading, and reinforced-but not opened-by PF binding.

## Feedback-primed accessible enhancers as a distinct regulatory class

The autoregulatory enhancers identified here not only drive expression of mesodermal determinants such as *odd* and *sna*, but also maintain accessibility through self-sustaining feedback loops. Their function does not require external opening by classic pioneer factors, placing them in a distinct regulatory category we term “feedback-primed accessible enhancers”. These elements act as regulatory landmarks that are preconfigured to be accessible and competent, even in chromatin environments enriched for Polycomb-associated repressive marks.

This autonomous behavior is further supported by the presence of H3K27me3 at active and accessible autoregulatory enhancers. Traditionally viewed as a silencing mark, the enrichment of H3K27me3 at these sites, absent from the Opa-dependent enhancers, suggests a modulatory rather than strictly repressive role. This aligns with recent studies showing that Polycomb complexes can coexist with transcriptionally permissive states, tuning enhancer activity rather than fully repressing it^53^.

Our data challenge the conventional binary view of pioneer factors as either gatekeepers or dispensable regulators. Instead, we suggest a more nuanced role for these factors, including functions in enhancer fine-tuning, stabilization, or burst regulation. Their binding at already-accessible enhancers may reflect enhancer bookmarking or support a poised chromatin state, rather than direct remodeling activity.

This model resonates with work from the Blythe group, which showed that Zld functions not only to open chromatin but also to buffer against Polycomb silencing at specific promoter-proximal elements ^53^. Taken together, our findings and theirs suggest that pioneer factors may be bifunctional: actively antagonizing repression at some loci, while acting as non-essential participants at others where accessibility is maintained through transcriptional feedback or architectural features.

### Polycomb regulation and enhancer poising

Autoregulatory enhancers in the mesoderm appear to reside in a poised or “ready” state, where Polycomb-mediated repression is present but not dominant. The overlap of Su(H) binding with H3K27me3-enriched domains further supports a model in which enhancers are held in check, yet primed for rapid activation. Su(H), as a Notch effector, parks at Polycomb-permissive enhancers and awaits NICD-mediated activation, consistent with poised enhancer models in both *Drosophila* and mammals.

These enhancers bear many hallmarks of developmental high occupancy regions: they are TF-rich^56^, accessible and positioned near promoters. While H3K27me3 is often interpreted as a repressive mark, our results, and those of others, suggest it may instead act as a tuning mechanism, preserving enhancer readiness while preventing premature activation. This poised configuration allows for fast, signal-dependent transcriptional activation, as might occur during gastrulation or lineage commitment.

Further supporting this poised model, we find evidence for paused RNAPII at many autoregulatory enhancers, even under transcriptionally arrested conditions. This suggests that enhancer accessibility can persist independently of active transcription, potentially maintained through RNAPII pausing and TF bookmarking. This finding mirrors observations from our single-nucleus multiomic data, in which chromatin accessibility at enhancers remains a more robust predictor of cell type than promoter accessibility, reinforcing the primacy of enhancer regulation in early embryonic patterning.

CBP, a key chromatin coactivator, plays a central role in stabilizing enhancer activity. Its maternal depletion leads to loss of transcriptional activity and cellular identity, yet accessibility at enhancers persists. This reveals a two-step model for enhancer activation: initial accessibility is established through pioneer or autoregulatory mechanisms, followed by CBP-mediated stabilization and activation of transcription.

### Evolutionary and functional implications

The existence of autoregulatory, pioneer-independent enhancers suggests a conserved strategy for stabilizing cell identity through intrinsic regulatory feedback. Analogous elements have been identified in vertebrate systems, for instance, the autoregulatory intronic enhancer of *NKX2-5* in cardiac mesoderm^57^, which further underscores the evolutionary relevance of these mechanisms. Feedback-primed accessible enhancers may therefore represent a foundational unit of developmental regulation, capable of both initiating and preserving transcriptional states across species.

Together, our findings reframe the canonical view of enhancer activation and chromatin accessibility by establishing that some enhancers function autonomously, independent of traditional pioneer factor input. These enhancers maintain accessibility and instruct cell identity through transcriptional feedback, even within repressive chromatin contexts. The identification of these “mentors of the genome” invites a re-evaluation of enhancer logic in early development and provides a conceptual framework for understanding lineage-specific gene regulation that is resilient, modular, and intrinsically encoded.

Future studies should focus on dissecting the molecular mechanisms underpinning feedback-mediated accessibility, the identity of potential co-factors that stabilize such enhancer states, and the extent to which this regulatory architecture is conserved across developmental systems and organisms.

Group IIB enhancers span genes expressed not only in mesoderm but also in neuroectoderm and endoderm, revealing a regulatory logic that cuts across lineage boundaries. These enhancers are accessible from very early nuclear cycles-well before cellularization-despite many of the associated genes not being transcriptionally active at those stages. This suggests that chromatin accessibility at these loci is not instructive per se, but permissive, setting up a regulatory state poised for future activation. In these cases, PFs such as Opa and GAF do not act as classical “openers” of chromatin but instead serve a maintenance or mentoring role, reinforcing enhancer activation and ensuring transcriptional fidelity during transitions in developmental state.

Our findings support a more nuanced view of PF function: not all PF binding events are equal, nor do they universally induce chromatin opening. Instead, PFs act within a hierarchical enhancer architecture, where their regulatory roles depend on both genomic context (e.g., proximity to promoters, pre-bound Pol II) and developmental timing. At distal group I enhancers, PFs are essential to initiate accessibility and transcriptional activation, often overcoming repressive chromatin environments marked by H3K27me3. In contrast, at proximal group IIB enhancers, PFs appear to stabilize already open regions, possibly acting as redundancy buffers or as part of a broader feedback loop that maintains gene expression robustness.

Importantly, group IIB enhancers are enriched near genes involved in lineage specification and tissue morphogenesis, particularly within mesodermal and neuroectodermal lineages, suggesting that these pre-accessible enhancers represent developmental “bookmarks” that prime cells for rapid transcriptional activation during later stages. Their accessibility and transcriptional readiness may reflect a conserved regulatory mechanism that balances developmental plasticity with precision, allowing cells to remain competent for multiple fates until lineage commitment solidifies.

To sum up, this enhancer architecture reflects a flexible yet robust framework for gene regulation during early embryogenesis, enabling the embryo to coordinate widespread gene activation while maintaining tight control over spatial and temporal expression patterns. These findings not only deepen our understanding of developmental enhancer logic but also inform broader models of cis-regulatory function, pioneer factor biology, and chromatin-based cell fate control.

### Materials and Methods

#### Fly stock and husbandry

*yw* strain (Bloomington *Drosophila* Stock Center (BDSC) #1495) embryos were collected as a wildtype control. To produce homogeneous populations of knockdown embryos, we employed RNA interference *(RNAi)* to knockdown the *opa* and *Su(H)*. *UAS-OpaRNAi* (BDSC #34707) and *shRNAiSu(H)* (BDSC #28900) females were crossed to Tubulin-Gal4 (Tub-Gal4) males (BDSC #5138), F1 female virgins were crossed back to *UAS-OpaRNAi* males to achieve the knockdown embryos. F2 embryos were collected from the apple agar plate at 26.5°C.

#### Fluorescent hybridization and Imaging

Following the meta-analysis, we designed a FISH experiment to examine the expression of *odd* and *slp1* genes in different genetic backgrounds. For controls of the experiment, we used wild-type (*yw*) and UAS-Tub-Gal4 embryos. Additionally to knockdown the expression of *opa* and *Su(H)* we used shRNAi (*sh-opa* and *sh-Su(H)*). We crossed this *sh_opa* and *sh_Su(H)* lines with Tub-Gal4 flies, to drive the expression in the early embryo. UAS-Tub-Gal4 and *OpaRNAi-Tub.Gal4* and *sh_Su(H).TubGal4* embryos were collected over a 2-4hour period and then processed using established methods for fixation and staining, with all procedures carried out at a constant temperature of 23°C. All specimens were simultaneously gathered, stained, and processed. Subsequently, they were imaged using Zeiss LSM 900 confocal microscope using a 20x air lens using 488 nm, 561 nm, and 647 nm lasers, with the imaging parameters kept consistent across all samples. Fiji (ImageJ) was used to process the obtained images^58^. fluorescent hybridization were performed with antisense RNA probes labeled with digoxigenin and biotin to visualize in vivo gene expression. *odd*, *slp1*, and *opa* riboprobes were used for FISH.

#### ChIP-seq and multiomics meta-analysis

To determine each TF binding profile, we used publicly available ChIP-seq data for TFs Dl at 2-4 hour, Twi at 2-4 hour, Opa 2-4 hour, GAF 2-4 hour, and Su(H) at 2-4, Zld, E(z), CBP, and RNA pol II, and they are aligned the libraries on the UCSC Genome Browser platform from the University of California, Santa Cruz^11,23,49,53,59–62^. To analyze the *in vivo* binding sites of Twi, Opa and Su(H), we utilized available ChIP-seq data from previously published datasets^11,23,29^. Peak calling was performed for Twi, Opa and Su(H) datasets^11,23,29^ to identify enriched regions corresponding to GAF, Dl, Twi, Opa and Su(H) binding sites across the genome. The data were processed using the MACS algorithm to refine the peak identification and assess the statistical significance of the binding events. This analysis allowed for the precise localization of Opa and Su(H) binding regions, which were further examined for overlap with known regulatory elements and target genes.

The overlapping regions of these TFs visualized using VennDiagram R package ChIPpeakAnno. To distinguish promoter-associated from non-promoter peaks, ChIP-seq binding sites were annotated relative to transcription start sites (TSS) using the *D. melanogaster* TxDb annotation (TxDb.Dmelanogaster.UCSC.dm6.ensGene) package. Peaks located within –500 to +100 bp of a TSS were classified as promoter peaks, whereas all other peaks were designated as non-promoter peaks. To visualize the overlap patterns among Dl, Twi, Opa, GAF, Su(H) and Zld ChIP-seq bindings, UpSet plots were generated using Python (pandas and the upsetplot library). The UpSet plots were generated by grouping ChIP-seq peaks according to the combination of TFs detected at each peak location. For promoter and enhancer-specific plots, peaks were classified as promoter-associated if they overlapped the region from –500 bp upstream to +100 bp downstream of a gene’s TSS, all remaining peaks were classified as enhancers. To avoid artificial inflation of counts, duplicate peak entries arising from associations with multiple regulatory elements were removed prior to visualization. UpSet plots were produced with upsetplot and matplotlib using default parameters and a custom TF order for clarity. To explore the histone marks modifications, we used H3K27ac and H3K27me3 CUT&Tag data^49^. Regions were processed in R as genomic ranges, centered on the peak summits and extended ±500 bp, and signal matrices were computed for each region. Heatmaps were produced using the EnrichedHeatmap and ComplexHeatmap packages, with regions stratified by group. Key regulatory elements were highlighted via row annotations. To investigate differential histone binding across mesodermal clusters, we used H3K27me3 and H3K27ac CUT&Tag data to generate histone mark profiles corresponding to mesodermal clusters identified from publicly available single-cell RNA-seq data^30^.

We conducted a meta-analysis ATAC-seq to investigate the chromatin accessibility landscape under six distinct conditions: ectopic expression of *opa*, and knockdown of *opa*, GAF and Dl knockdown, Zld mutant, and wild-type control for different timepoints at nc14^11,26,60^.

Stage 5 specific RNA-seq data^55^ were used to compare the expression levels of *odd*, *run*, *prd, sna* and *slp1*. Violinplot was created using this dataset and a gray gradient was used for violin fills to clearly distinguish the timepoints while maintaining visual clarity. Normalized gene expression values for individual embryos at nuclear cycle stages nc14A-D were plotted using violin plots to show the distribution of expression for each gene. Individual replicate points were overlaid using jittered scatter points to display variability across embryos. Pairwise comparisons between selected timepoints (nc14A vs nc14C, nc14A vs nc14D, nc14C vs nc14D) were performed using two-tailed t-tests, and statistically significant differences were indicated directly on the plots with asterisks (*p < 0.05, *p < 0.01, **p < 0.001). Plots were generated in R (version R 4.5.0) using the ggplot2 and ggpubr packages. A gray gradient was used for violin fills to clearly distinguish the timepoints while maintaining visual clarity. Each gene was shown in a separate facet to allow easy comparison of expression patterns across timepoints.

Gene Ontology (GO) enrichment analysis was performed for both group I and group II using the clusterProfiler R package. Only terms with a false discovery rate (q-value) < 0.05 were considered significant. The top enriched GO categories were visualized as bar plots, showing −log10(q-value), generated with ggplot2.

The single-cell data that were publicly available^30^ were analyzed using the Scanpy standard pipeline with default parameters (resolution = 0.9). Subclustering and cell type annotation was guided by marker genes reported in the supplementary of ref. 30 and spatial information using DistMap^30^. Candidate marker genes were defined by differential expression analysis using Wilcoxon rank-sum tests. To increase robustness, we required markers to satisfy thresholds on fold-change and detection rates. Specifically, for pairwise comparisons, each cluster was compared to its two nearest neighbors in PCA space, and genes were retained if they met the following criteria in both comparisons: maximum out-group fraction ≤ 0.5, log fold-change ≥ 0.5, and in-group fraction ≥ 0.5. For global comparisons, each cluster was compared against all remaining clusters, with stricter thresholds (out-group fraction ≤ 0.5, log fold-change ≥ 0.5, in-group fraction ≥ 0.7). The intersection of pairwise and global marker sets was used as the final marker set for each cluster.

For bulk RNA-seq data processing, paired-end fastq sequencing data from GSE153328 were processed using rsem-calculate-expression with Bowtie2 for alignment. Only replicates 1 and 3 from both control and sh_opa were used in downstream analysis. After filtering, only genes with TPM > 10 in at least two samples (since there were two replicates per condition) were retained. For mesoderm-specific genes, a stricter threshold of TPM > 7 in at least two samples was applied. One-way hierarchical clustering was performed using Pearson correlation coefficients and average linkage on the expression profiles of genes under the two conditions.

## Acknowledgements

We would like to thank Mike Levine and Angela Stathopoulos for generously providing fly lines. Our gratitude extends to Marion Ball, Christos Delidakis, George Mosialos, Konstantinos Chatzistergos and Nikoleta Psatha for their insightful discussions and contributions. We further acknowledge all members of the Luber Lab for their productive input, and Rie Conley, Oscar Villasana Espinosa, Georgios Tegousis, and the Koromila Lab for their assistance with administrative tasks and fly husbandry. This research was supported through funding from the UTA STARS program.

