## Supplementary Figures for "Balancing precision with plasticity: Redefining the roles of transcription factors in early cell fate specification"

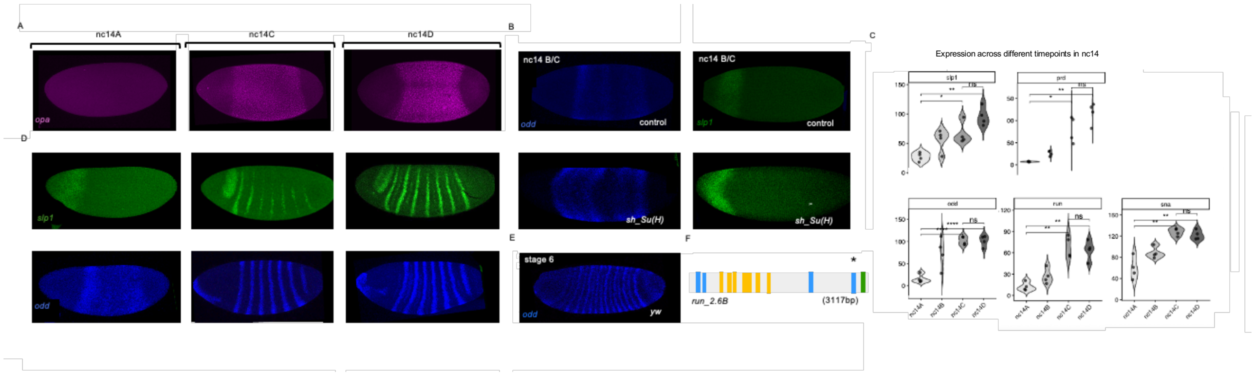


**Supplementary fig. 1.1 Regulatory dynamics and gene expression changes during early embryogenesis** (A, D) in situ hybridization of embryos at nc14A, mid-nc14C, and nc14D demonstrating the dynamic expression patterns of Opa (magenta), *odd* (blue), and *slp1* (green) at indicated stages. The embryos are positioned with the anterior facing left and the dorsal side facing upward.

B)Su(H) roles in regulating *odd* and *slp1*. (A) *Su(H)* knockdown results to alteration in *odd* and *slp1* expression at cellularization. *In situ* hybridization of *odd* (blue) and *slp1(*green) in control (UAS-lacZ.Tub-Gal4) embryos and *sh_Su(H)*.

C) RNA-seq expression level comparisons for *odd*, *run*t (*run*) (two primary pair-rule genes) and *slp1* and *paired* (*prd*) (two secondary pair-rule gene) at nuclear cycles 14A-14D. All figures were created using the RNA-seq publicly available data. Statistical comparisons between selected stages highlight significant differences (*p < 0.05, **p < 0.01, ***p < 0.001). Separate panels display each gene for cross-stage comparison.

E) In situ hybridization for *odd* in a *yw* stage 6 embryo.

F) Schematic showing transcription factor binding sites for Dl, Opa, Su(H), and GAF within the genomic region of *run_2.6B* (3117 bp). See key in fig. 1G.

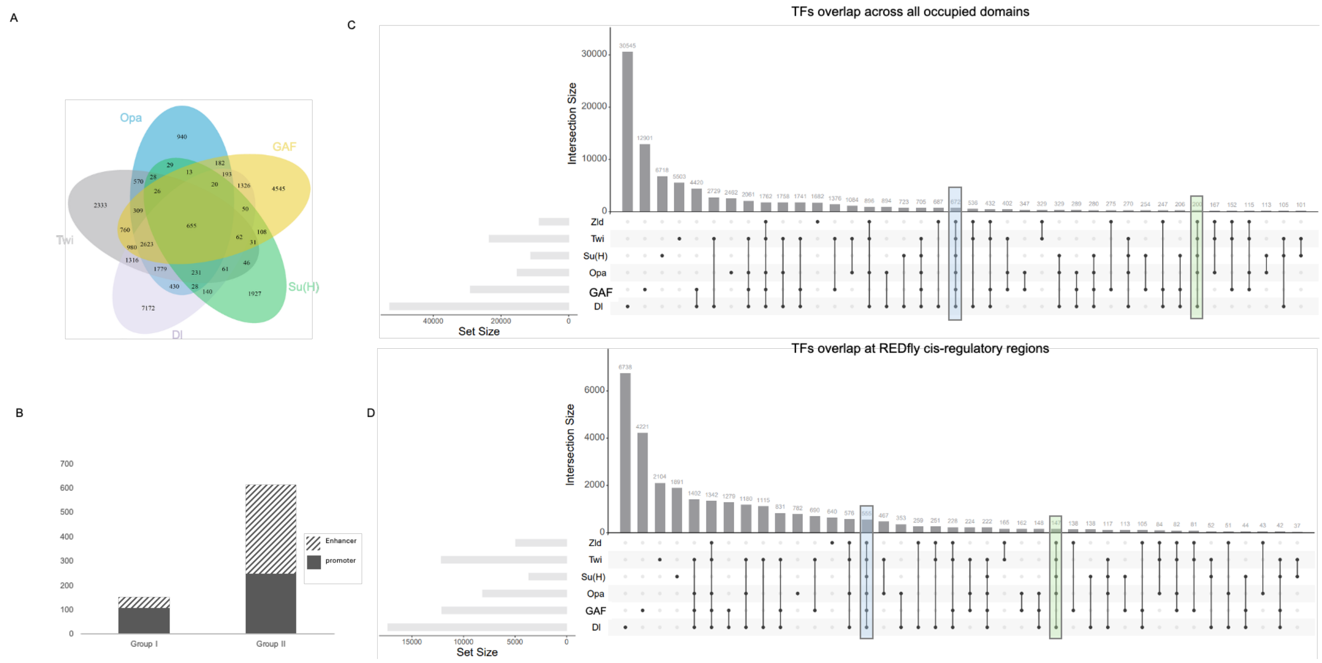


**Transcription factor binding patterns in *Drosophila* embryos**. A) Venn diagram generated using publicly available ChIP-seq data, showing overlapping and unique binding sites among Twi (gray), Opa (blue), Dl (Lavender), GAF (yellow), and Su(H) (green)[11,23,29].TF peaks were filtered to retain only those overlapping within 100 bp of REDfly cis-regulatory modules. B) Bar plots showing the proportion of regions located in promoters and enhancers within group I and group II.

C) Upset plot showing overlap of Dl, GAF, Zld, Twi, Opa, and Su(H) TFs at known cis-regulatory regions. Overlap located at regulatory regions or within 100bp of an enhancer (Bottom) and all the peaks (Top). Bars indicate the number of regions in each intersection, with combinations represented by filled circles below. Group I and group II categories are highlighted with green and blue boxes, respectively.

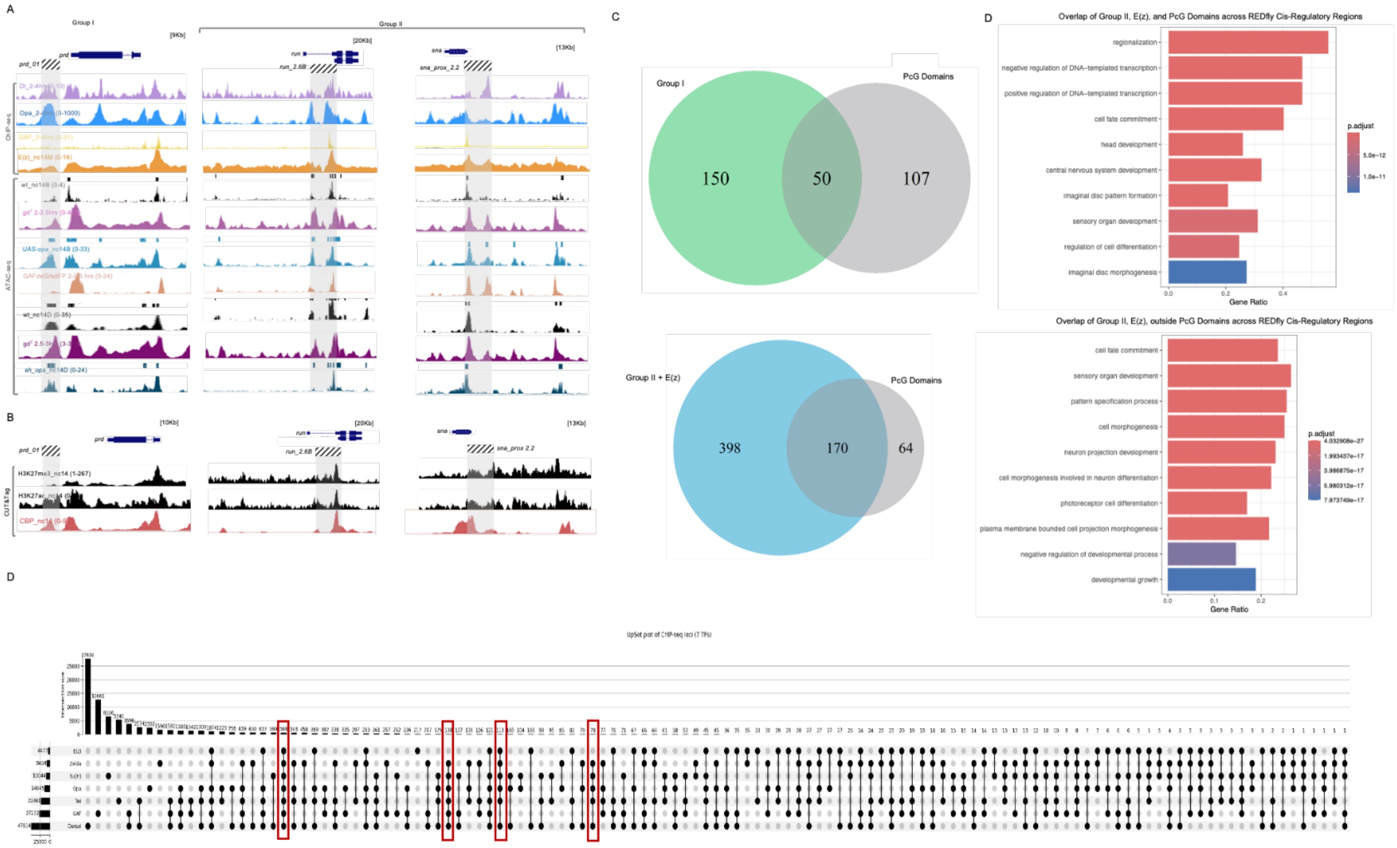


**Supplementary fig. 2.1. TFs Bindings and chromatin accessibility profiling.** A) ChIP-seq of Dl at 2-4hrs (lavender), Twi at 2-4hrs (gray), Opa 2-4hrs (blue) and Su(H) at 2-4hrs (green), and GAF (in yellow) from available data[11,23,28,29] in *prd* (groupI) and *run, sna* (group II) ATAC-seq tracks represent wildtype control for nc14B (wt) (top black), *UAS-opa*_nc14B (light blue), wt control for nc14D (bottom black), *sh_opa* knockdown at nc14D (dark blue). GAF degraded embryos 2–2.5 hrs (yellow), Dl mutant/embryos lacking nuclear Dorsal 2-2.5 hrs (light purple) and gd$^7$/embryos lacking nuclear Dorsal 2.5-3 hrs (dark purple) in *prd_01, run 2.6B, sna_prox 2.2* and *odd_basal* loci.All the regulatory elements are highlighted in gray. B) Analysis of H3K27me3, H3K27ac, and CBP occupancy at the prd_01, run_2.6B, sna_prox 2.2 luci during nc14. C) Upset plot showing the overlap of seven transcription factors. Overlap located at regulatory regions or within 100bp of an enhancer (Bottom) and all the peaks (Top). Bars indicate the number of regions in each intersection, with combinations represented by filled circles below. Group I and group II categories are highlighted with green and blue boxes, respectively. D) Visualization of GO term enrichment among genes classified into group II and PcG domain overlap and group II non-PcG domain in REDfly Cis-Regulatory regions, highlighting functional differences between the two groups.

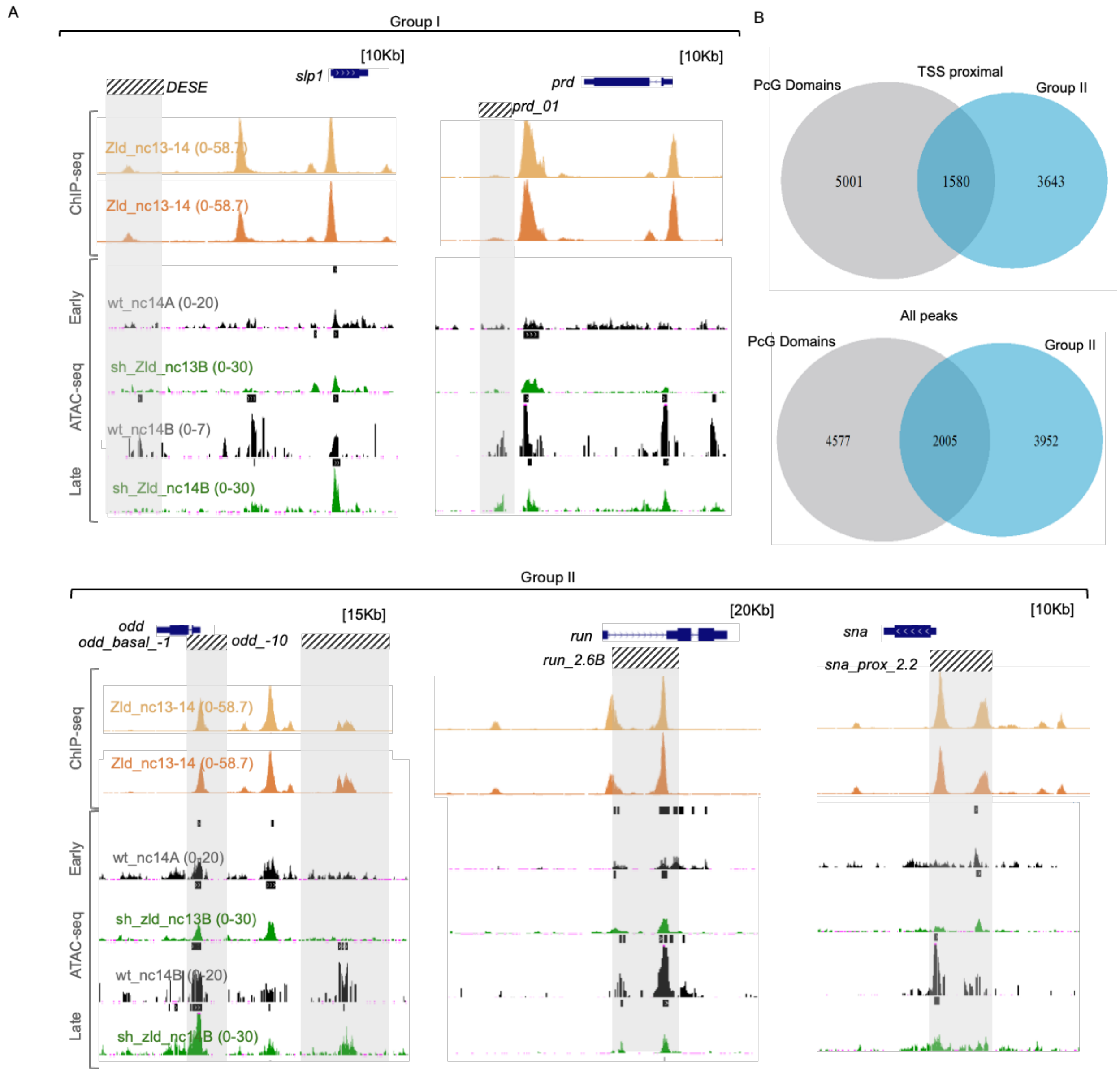


**Supplementary fig 2.2. Zld binding and chromatin accessibility in enhancer groups, with PcG domain association in group II**. A) UCSC browser tracks displaying Zld ChIP-seq at nc13–14 (light orange) and nc14 (dark orange), alongside ATAC-seq profiles from wild-type (nc14A, nc14B) and *sh_zld* (nc13B, nc14B) embryos obtained from available datasets. B) Overlap between group II and PcG domains near TSS (top) and across all peaks (bottom).

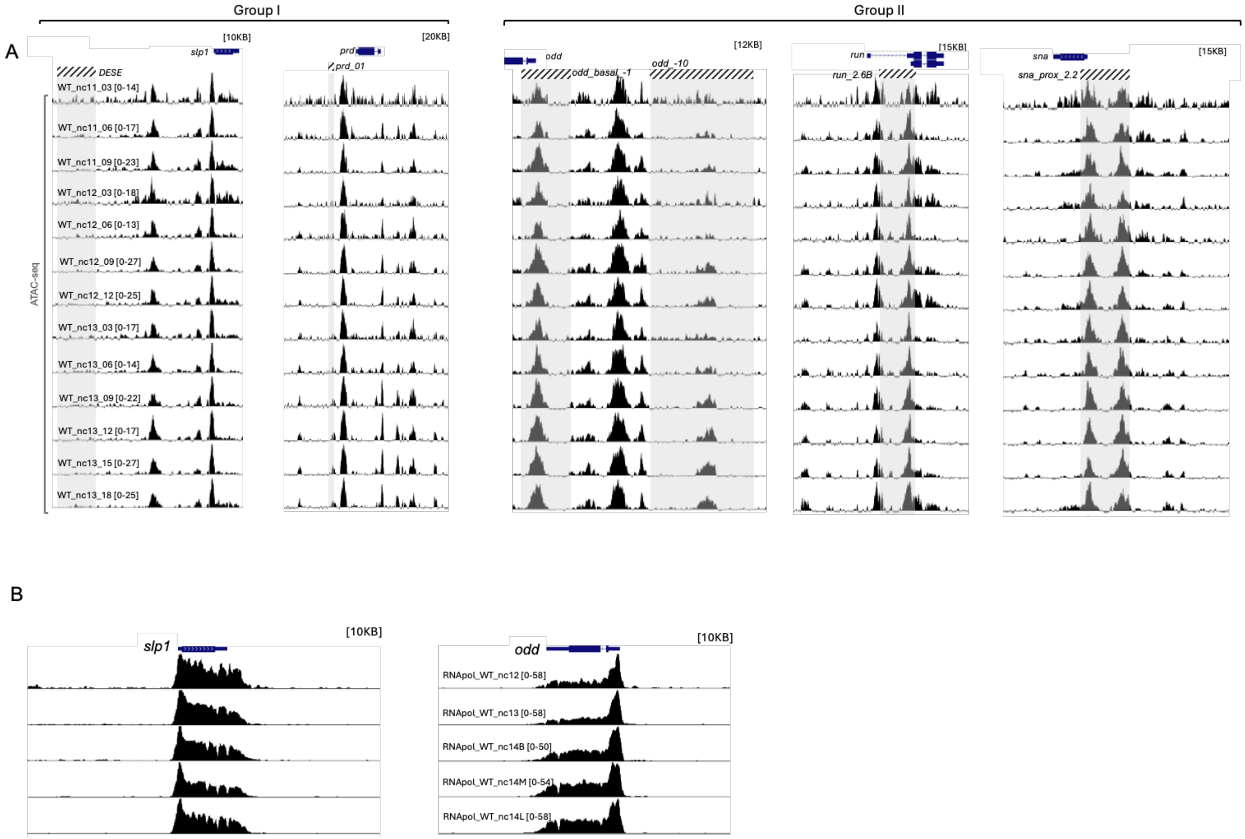


**Supplementary fig 2.3. Chromatin accessibility and RNApol II activity within two groups.** (A) Chromatin accessibility of enhancers from groups I and II. Tracks display ATACseq profiles from wild-type embryos collected at 3-minute intervals, spanning nc11 to late nc13. (B) RNA Pol II occupancy near the TSS of *odd* and *slp1*.

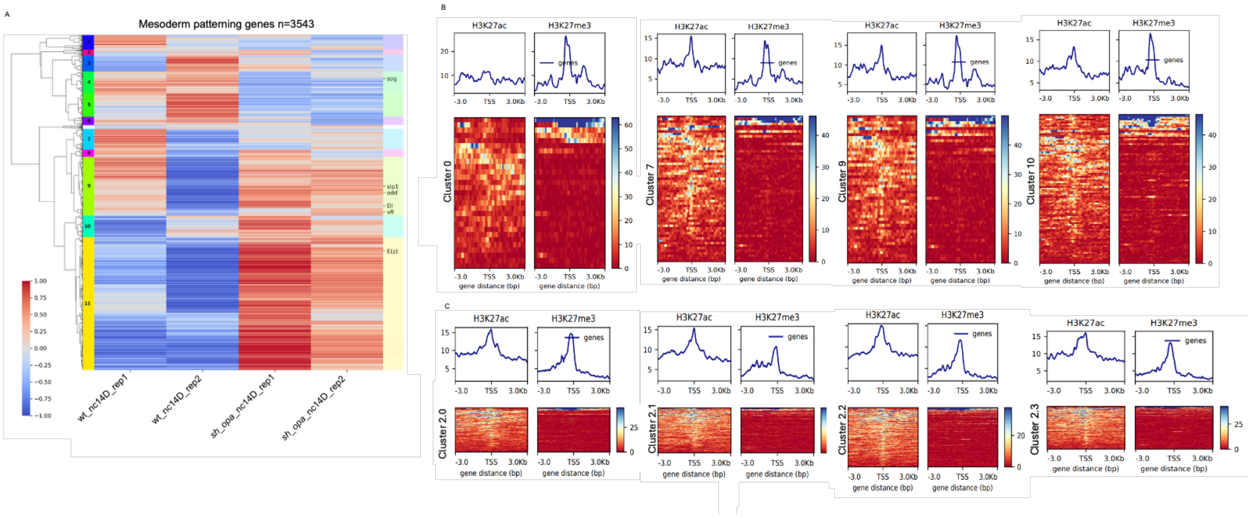


**Supplementary fig. 3.1. Mesodermal gene clusters and chromatin accessibility among different mesodermal clusters.** A) Heatmap of mesoderm patterning genes, defined as genes from mesodermal cluster genes heatmap (fig. 3C) after subtracting mesoderm-specific genes. B,C) TSS-centered heatmaps showing the occupancy of histone marks H3K27ac and H3K27me3 across clusters 0, 2, 7, 9 (B), and subclusters 2.0, 2.1 and 2.2 (C ). All heatmaps were generated using available CUT&Tag data and by deepTools.

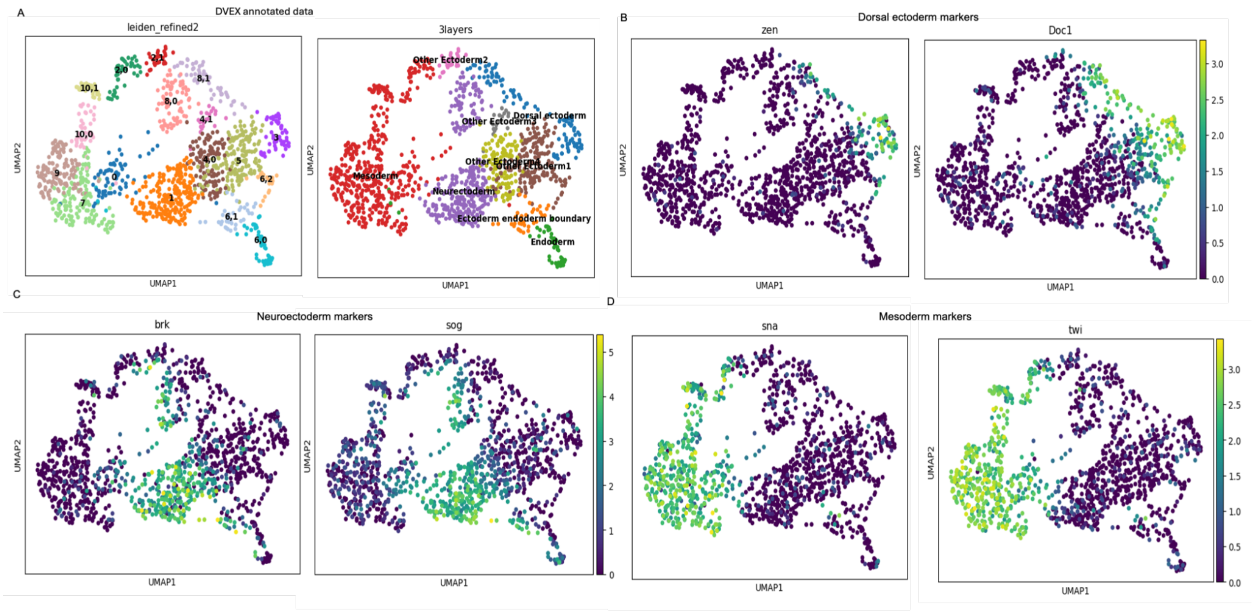

**Supplementary Fig. 3.2. UMAP visualization of germ layer identity.** (A) UMAP plot showing distinct cell clusters and their corresponding germ layers (DVEX annotated data) (B–D) UMAP plots of representative germ layer markers: dorsal ectoderm (*zen*, *Doc1*), neuroectoderm (*brk*, *sog*), and mesoderm (*sna*, *twi*). The key shows he color scale indicates relative gene expression levels across individual cells.

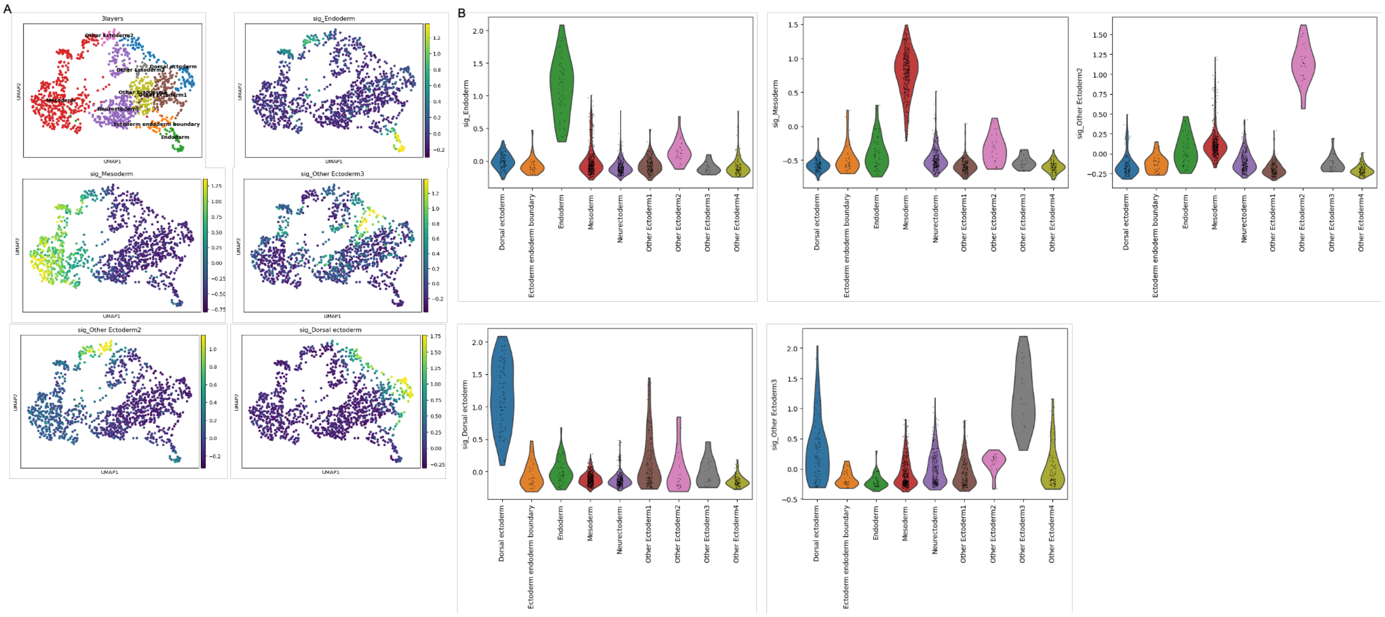


**Supplementary fig. 3. 3. Identification of germ layers using UMAP and marker gene expression**

A) UMAP plot showing the major germ layers. Subsequent UMAPs display the expression patterns of individual germ layer markers. B) Violin plots showing the expression distribution of germ layer marker genes across the corresponding germ layers.
